# Mitochondrial transplantation: A novel therapy for liver ischemia/reperfusion injury

**DOI:** 10.1101/2024.09.04.608457

**Authors:** Avinash Naraiah Mukkala, Bruna Araujo David, Menachem Ailenberg, Jady Liang, Chirag Manoj Vaswani, Danielle Karakas, Rachel Goldfarb, William Barbour, Avishai Gasner, Ruoxian Scarlet Wu, Raluca Petrut, Mirjana Jerkic, Ana C. Andreazza, Claudia dos Santos, Heyu Ni, Haibo Zhang, Andras Kapus, Paul Kubes, Ori David Rotstein

## Abstract

Mitochondrial transplantation prevented liver ischemia/reperfusion-induced hepatocellular injury and inflammation. *In vivo* intravital microscopy demonstrated that liver resident macrophages, namely Kupffer cells, rapidly sequestered, internalized and acidified transplanted mitochondria through the CRIg immunoreceptor. Mechanistically, both Kupffer cells and CRIg were necessary for the hepatoprotective and anti-inflammatory effects of mitochondrial transplantation.

**STRUCTURED ABSTRACT:** *Objective:* To investigate the hepatoprotective effects of mitochondrial transplantation in a murine liver ischemia/reperfusion (I/R) model.

*Summary background data:* Sequential liver ischemia followed by reperfusion (I/R) is a pathophysiological process underlying hepatocellular injury in a number of clinical contexts, such as hemorrhagic shock/resuscitation, major elective liver surgery and organ transplantation. A unifying pathogenic consequence of I/R is mitochondrial dysfunction. Restoration of mitochondria via transplantation (MTx) has emerged as potential therapeutic in I/R. However, its role in liver I/R and its mechanisms of action remain poorly defined.

*Methods:* We investigated the hepatoprotective effects of MTx in an *in vivo* mouse model of liver I/R and used *in vivo* imaging and various knockout and transgenic mouse models to determine the mechanism of protection.

*Results:* We found that I/R-induced hepatocellular injury was prevented by MTx, as measured by plasma ALT, AST and liver histology. Additionally, I/R-induced pro-inflammatory cytokine release (IL-6, TNFα) was dampened by MTx, and anti-inflammatory IL-10 was enhanced. Moreover, MTx lowered neutrophil infiltration into both the liver sinusoids and lung BALF, suggesting a local and distant reduction in inflammation. Using *in vivo* intravital imaging, we found that I/R-subjected Kupffer cells (KCs), rapidly sequestered transplanted mitochondria, and acidified mitochondria within lysosomal compartments. To specifically interrogate the role of KCs, we depleted KCs using the diphtheria toxin-inducible Clec4f/iDTR transgenic mouse, then induced I/R, and discovered that KCs are necessary for the beneficial effects of MTx. Finally, we induced I/R in complement receptor of the immunoglobulin superfamily (CRIg) knockout mice and found that CRIg was required for mitochondria capture by KCs and mitochondrial-mediated hepatoprotection.

*Conclusions:* In this study, we demonstrated that CRIg-dependent capture of mitochondria by I/R-subjected Kupffer cells is a hepatoprotective mechanism *in vivo*. These data progress knowledge on the mechanisms of MTx and opens new avenues for clinical translation.

## INTRODUCTION

Ischemia/reperfusion (I/R) of the liver is a common pathophysiological process contributing to hepatocellular injury in the clinical settings of traumatic hemorrhagic shock, major liver operations and in liver transplantation.^1–3^ During the ischemia phase, the liver is deprived of blood flow, which depletes ATP, oxygen, and nutrient exchange, then in the reperfusion phase, the flow of oxygenated blood induces several cellular events that are deleterious to the cell.^4,5^ One unifying hallmark of I/R is the induction of mitochondrial dysfunction.^4,5^ The perturbation of the mitochondrial membrane potential produces reactive oxygen and nitrogen species, creating a positive feedback loop of oxidative stress.^4,5^ This contributes to direct cellular injury but also to the initiation of inflammatory signalling, activation of the innate immune system with neutrophil infiltration, macrophage activation and cytokine release.^1,6^

Given the central role of mitochondrial dysfunction in the pathogenesis of I/R, a novel therapeutic has emerged to target and correct mitochondrial injury by the introduction of exogenous mitochondria to the injured organ.^7^ This biologic therapeutic is termed ‘mitochondrial transplantation (MTx),’ and involves the injection/infusion of healthy, intact, and functional mitochondria into the damaged organ.^7^ MTx has been shown to be safe and efficacious in animal models of heart, lung, kidney, and brain I/R.^8–13^ Few reports have described the ability of MTx to lessen liver I/R,^8,9^ and these studies have been limited by failure to study mechanisms of action.

The mechanisms leading to the initiation of injury in response to liver I/R have been extensively studied. Kupffer cells (KCs) are liver tissue-resident macrophages situated in the sinusoids,^14^ and are known to mediate early injury through the propagation of free radicals, pro-inflammatory cytokines, and neutrophil activation.^14–16^ Together these lead to cellular injury resulting in hepatocellular necrosis and apoptosis and endothelial cell dysfunction causing microvascular injury. In addition, liver I/R causes systemic inflammatory response syndrome through KC activation, thereby expounding remote injury.^17^

In this article, we provide clear evidence that mitochondria delivered into the portal vein are rapidly captured by KCs in the liver leading to an immunophenotypic switch from a pro-inflammatory to an anti-inflammatory microenvironment and culminating in protection against hepatocellular injury as well as reducing systemic inflammation, including in the lungs. The protection occurred when mitochondria were injected following the ischemic phase and at the beginning of reperfusion, therefore supporting its potential for clinical application.

## METHODS

### Mouse liver I/R model

Mice (11±2w; 27±3g) were acclimated for 1 week. Partial 70% warm left/median lobe ischemia was achieved as described previously.^18,19^ Briefly, the left hepatic artery was clamped, to induce 1h of left/median lobe ischemia, confirmed by blanching of the lobes, while the right lobe remained perfused (non-ischemic). 1h of ischemia/2h of reperfusion was used in studies, except where indicated. Sham-operated mice underwent surgical manipulations except clamping. Ischemia-only mice underwent 1h of ischemia followed by sacrifice. Mice were exsanguinated, and blood and liver was collected.

### Animal ethics and mice

Animal studies were in compliance with the Animal Care Committee (ACC) at St. Michael’s Hospital (ACC #171) and the ACC at University of Calgary (AC19-0138). Wild-type (WT), male, C57BL/6J mice were acquired at 8-weeks old (JAX stock #000664). Complement receptor of immunoglobulin family knockout (CRIg^-/-^) (gene name: *VSIG4*) mice were obtained from Genentech. CRIg^-/-^ mice contain an exon 1 replacement which results in loss of the full-length functional CRIg protein, this was confirmed by Helmy *et al*.^20^ Transgenic Clec4f/iDTR mice were provided by Dr. Heyu Ni (JAX stock #007900 and # 033296).^21,22^ All mice were on a C57BL/6 genetic background. All mice were genotyped (Fig. S5).

### Mitochondria isolation, dose and transplantation

As per Preble *et al*., mitochondria were isolated from fresh hindlimb mouse skeletal muscle (MSM) tissue by differential filtration and centrifugation (protocol is detailed in reference).^23^ Mitochondria injected within 1-2h of isolation. Mitochondria were counted on a Coulter particle counter (Beckman), and 8x10^7^ mitochondria/mouse were intrasplenically injected.^9^ Mitochondria were treated with 50uM of antimycin-A (AntA) in respiration buffer (RB) and incubated on ice 30mins and washed 3 times to remove AntA. The vehicle control was RB. As a control for particle injection, 8x10^7^ polyethylene beads (PE, 1-4μm) were injected.

### Plasma ALT and AST

Fresh heparinized (∼2.5U/mL) mouse blood was centrifuged at 2500xg for 10mins at 4°C, and plasma collected. ALT and AST concentration was quantified within 24h by the St. Michael’s Hospital Diagnostic Laboratory.

### Liver histology, immunohistochemistry and quantification

Preparation of 5μm formalin-fixed, paraffin-embedded (FFPE) liver sections was described previously.^24^ FFPE slides were H&E stained. For immunohistochemistry, FFPE slides were stained for 4-hydroxynonenal (4-HNE) and detected with 3,3’-diaminobenzidine staining (Table S1). Images were acquired using the Zeiss Axio Scan (v2.1) system and analyzed in HALO (v2.3.2089.23). 3-5 independent blinded observers scored each mouse. The Suzuki Liver Injury Score was used for quantification of liver necrosis (0-5), hepatic congestion (0-4), and cytoplasmic vacuolization (0-4).^25^ The necrosis score was modified to 0-5, five indicating >80% confluent necrosis. For analysis of 4-HNE^+^ area, the HALO classifier was trained to identify positive and negative brown staining, and positive area was divided by total tissue area.

### Enzyme-linked immunosorbent assays

ELISA kits were used to measure the concentration of TNFα, IL-6, CCL2, 8-isoprostane, and IL-10 in plasma (Table S1). HGF was measured in liver homogenates (Table S1).

### ATP assays

ATP concentration in mitochondria pellets and liver tissue was measured using a luciferase assays (Table S1). Liver tissue ATP extraction was described previously.^26^ Modifications are in the supplement.

### Intravital confocal microscopy and quantification

Intravital microscopy was customised from Peiseler and David *et al*., Marques *et al.* and Surewaard *et al.*.^27–29^ Method and microscope parameters are in the supplement.

### Clodronate liposomes and diphtheria toxin Kupffer cell depletion

24h before surgery, transgenic Clec4f/iDTR mice were intraperitoneally injected with diphtheria toxin (DT, 10ng/g) or PBS control (Table S1). Clec4f/iDTR mice express the diphtheria toxin receptor (DTR) under direction of *Clec4f*, a lectin expressed on mouse KCs, upon IP injection of unnicked DT (derived from *C. diphtheriae*), KCs were selectively depleted.^21,22,30,31^ 48h before surgery, WT mice were tail-vein (IV) injected with 50ug/g clodronate liposomes (CL) to deplete macrophages (Table S1).^32^

### Study design and statistics

Injury and inflammation endpoints were blindly measured. Each mouse is considered one biological replicate, with assays in technical replicates. Statistical analyses were conducted using Prism (V10.1.1, GraphPad). For significance testing, ordinary one-way ANOVA with *post hoc* Tukey’s test was used, with *p<0.05 being considered statistically significant. Unpaired T-test was utilized when two groups were compared. Significance is noted with asterisks *p<0.05, **p<0.01, ***p<0.001 and ****p<0.0001.

### Other standard methods

Western blot, TEM, and qPCR were described previously,^33^ with modifications in the supplement. ddPCR,^34,35^ mitochondria nanocytometry,^36^ flow cytometry,^32^ and BALF^37,38^ were described previously, with modifications in the supplement.

## RESULTS

### Effect of mitochondrial isolation technique on mitochondrial structure and function

To assess the integrity of the mitochondrial membrane and cristae architecture, we employed TEM of isolated MSM mitochondria. Mitochondria had intact double-membrane and cristae architecture (Fig. 1A). To confirm mitochondrial proteins in our isolates, we stained mitochondria with ATPB-AlexaFluor647 antibody and identified positive labelling, compared to unstained control (Fig. 1B). To confirm enrichment of mitochondrial proteins in isolates, we performed Western blotting for TOM20 and ATPase F_1_F_0_ subunit (complex V), as representative of inner and outer mitochondrial membranes, respectively. Both TOM20 and ATPase F_1_F_0_ subunit were enriched in mitochondrial pellets, whereas cytoplasmic proteins, such as GAPDH, were diminished (Fig. 1C).

**Fig. 1.**
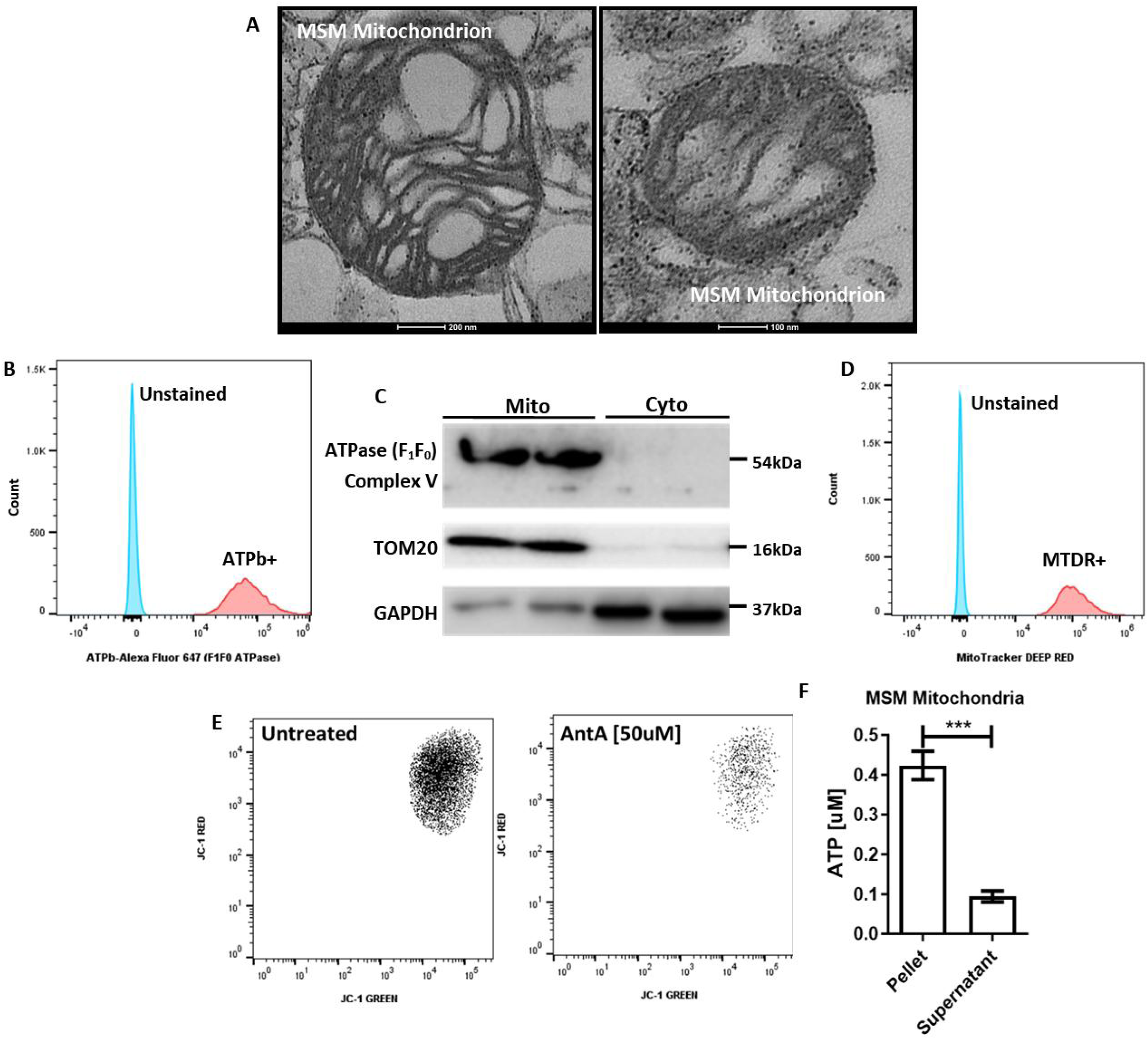
Effect of mitochondrial isolation procedure on mitochondrial structure and function. (**A**) Representative transmission electron microscopy images of individual isolated mitochondria from mouse hindlimb skeletal muscle tissue (n=3 independent isolations). (**B**) Abolishment of mitochondrial membranes and cristae architecture by liquid nitrogen free- thawing. (**C**) Nanocytometry of ATPB-conjugated-Alexa Fluor 647 labelled mitochondria showing positive labelling compared to the unstained control (n=2 independent isolations). (**D**) Western blots of isolated mitochondria showing enrichment of mitochondrial proteins TOM20 and ATPase F_1_F_0_ subunit, and reduction in cytoplasmic protein GAPDH (n=2 independent isolations). (**E**) Positive labelling of isolated mitochondria with the mitochondria-specific dye MitoTracker DEEP RED (MTDR). (**F**) Staining of isolated mitochondria with JC-1, a marker of mitochondrial membrane potential (ΔΨ_m_), showing reduced JC-1^+^ (red-green-positive) population after liquid nitrogen membrane disruption. (**G**) Treatment of isolated mitochondria with antimycin-A (50uM) diminished the JC-1^+^ population, indicating impairment of mitochondrial function (ΔΨ_m_). (**H**) Measurement of ATP content by luciferase bioluminescence assay determined that mitochondria pellets were enriched in ATP, and the cytoplasmic supernatants were depleted of ATP (n=3 independent isolations).

As a measure of mitochondrial functionality, isolates were stained for mitochondria-specific dyes, JC-1 is a mitochondria-selective dye for the binary measurement of mitochondrial membrane potential (ΔΨ_m_), and MitoTracker DEEP RED (MTDR) sequesters to functional mitochondria. Nanocytometry confirmed positive staining for both MTDR and JC-1, thereby confirming the ability of isolated mitochondria to maintain ΔΨ_m_ (Fig. 1D-E). By contrast, we treated isolated mitochondria with AntA (a specific complex III irreversible inhibitor) and showed that there was a diminution of ΔΨ_m_ (Fig. 1E). Finally, by luciferase assay, mitochondria pellets were significantly enriched in ATP, as compared to cytoplasmic supernatants (Fig. 1F). In aggregate, these experiments confirmed the structure, purity, and functionality of isolated MSM mitochondria for use in *in vivo* studies.

### Mitochondrial transplantation prevents I/R-induced hepatocellular injury

Mouse liver I/R injury was generated by 1h occlusion of blood flow into the left/median lobes, followed by release allowing reperfusion over time. We used intrasplenic injection of mitochondria (8x10^7^/mouse) which resulted in a rapid infusion into the portal vasculature within seconds of splenic injection (Movie S1). Injection of mitochondria began as the arterial clamp was removed, at t=0h reperfusion.

At t=0h, plasma hepatocellular enzyme levels were low, approximating that observed in sham-operated animals (Fig. 2A-B). Following reperfusion, there was a time-dependent increase in hepatocellular injury as measured by plasma ALT and AST levels (Fig. 2A-B), reaching a maximum at 2h. MTx significantly reduced hepatocellular enzymes compared to control groups, indicative of hepatocellular protection (Fig. 2A-B; blue bar 5). To determine whether an active mitochondrial electron transport chain was required for this protection, we pretreated isolated mitochondria with AntA. We found that injection of AntA-treated mitochondria was not hepatoprotective (Fig. 2A-B, bar 6). Neither injection of polyethylene (PE) beads (size range 1- 4um), to rule out physical effects of particle injection, nor injection of respiration buffer (RB), as a control for splenic injection, *per se*, were protective (Fig. 2A-B, bar 7 and 8). Together, these studies demonstrate that MTx reduces hepatocellular injury in a murine liver I/R injury and that functional mitochondria are required for this effect.

**Fig. 2.**
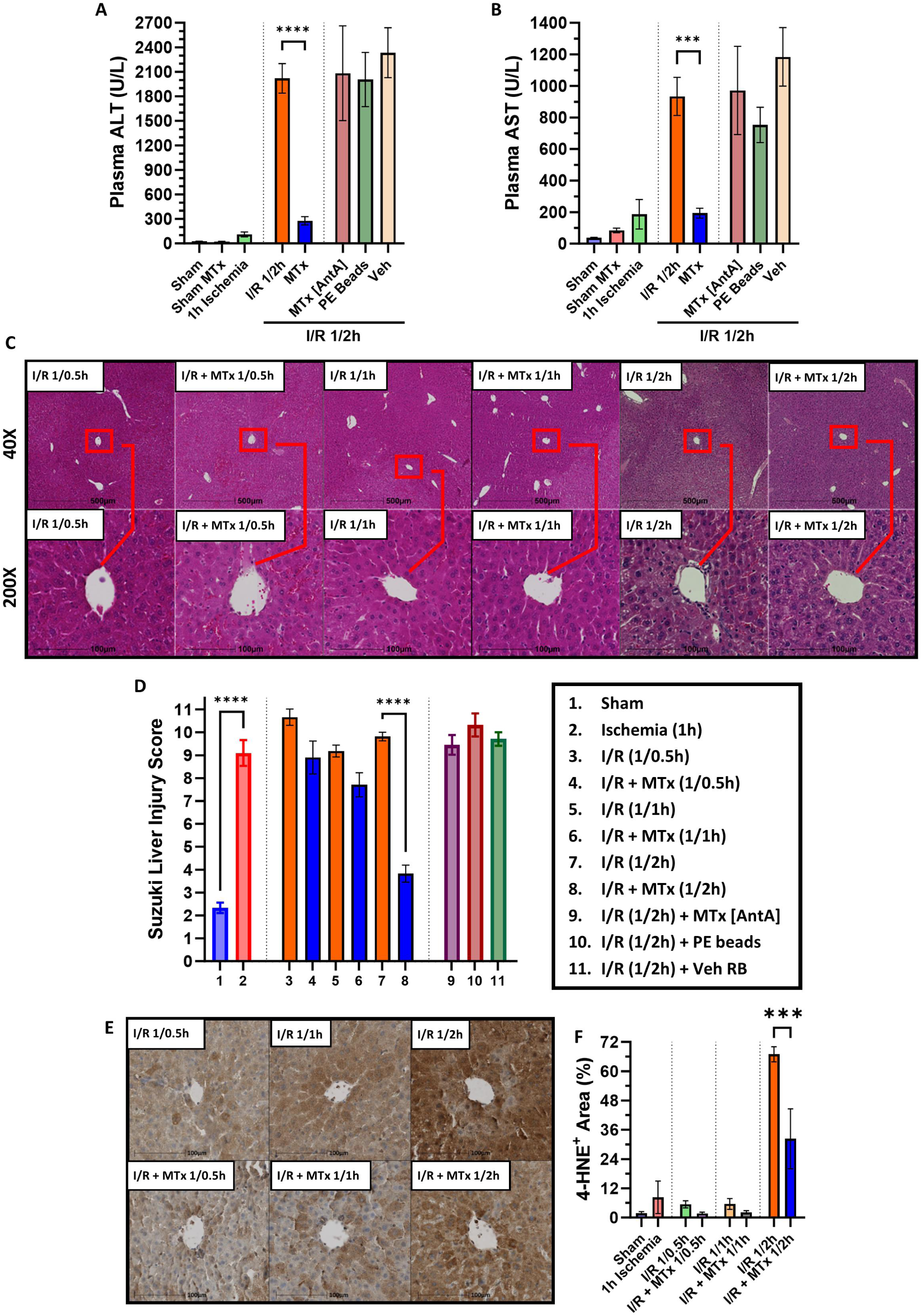
Mitochondrial transplantation prevents I/R-induced hepatocellular injury. (**A-B**) MTx significantly reduced plasma ALT and AST, following 1h ischemia and 2h reperfusion in WT C57BL/6J mice, demonstrating hepatoprotection (p<0.0001; n=13-26 mice in sham, I/R 1/2h and I/R + MTx 1/2h groups, n=3-10 mice in all other groups). (**C**) Representative time- course liver histology of liver I/R control and liver I/R + MTx mice. Liver I/R + MTx mouse liver sections exhibit lowered cytoplasmic vacuolization, hepatic congestion, and zone 2/3 confluent necrosis compared to liver I/R control mice. (**D**) Suzuki liver injury score quantified by 3-5 blinded observers per mouse. Liver I/R control mice showed no resolution of hepatocellular injury, whereas liver I/R + MTx mice showed a progressive resolution of injury (p<0.0001; n=10-28 mice in groups 1, 7, 8, n=4-10 mice in all other groups). (**E**) Representative liver 4- HNE immunohistochemistry sections of liver I/R control (top row) and liver I/R + MTx (bottom row) mice. With increasing reperfusion time, the liver I/R control mice have increasing 4-HNE lipid peroxidation, whereas liver I/R + MTx mice show significantly lowered 4-HNE staining at 2h of reperfusion. (**F**) Quantification of 4-HNE positive area percentage demonstrated that liver I/R + MTx mice had reduced lipid peroxidation (rightmost blue bar) (p<0.0001; n=4-6 mice in each group).

To better illustrate the effects of MTx on hepatocellular injury in liver I/R, we performed H&E histology (Fig. 2C) of mouse liver tissue and utilized the Suzuki score to blindly quantify hepatocellular injury (Fig. 2D). Ischemia-only (1h) significantly increased Suzuki score, compared to sham-operated mice (Fig. 2D). MTx at the beginning of reperfusion significantly lowered the Suzuki score, as compared to liver I/R control mice at 2h following reperfusion (Fig. 2C-D). To better understand the hepatoprotective effects of MTx in liver I/R, we performed a time-course of liver histological findings. Over the 2h of liver reperfusion, untreated I/R mice showed no resolution of hepatocellular injury, as quantified by the Suzuki score (Fig. 2C-D). By contrast, mice injected with mitochondria showed progressive resolution of histological changes, reaching significance by the 2h time point (Fig. 2C-D, bars 4,6,8). PE beads injected I/R mice and vehicle RB injected I/R mice (Fig. 2D, bars 10,11) did not show any resolution of Suzuki score. Finally, injection of AntA-treated mitochondria in I/R mice, did not attenuate the Suzuki score (Fig. 2D, bar 9). Therefore, the injection of intact and functional mitochondria promoted the progressive resolution of the histological injury associated with liver I/R. Sham mice showed no evidence of hepatocellular injury (Fig. 2D).

Oxidative stress induced by I/R contributes to hepatocellular injury. We measured tissue 4-hydroxynonenal (4-HNE) by immunohistochemistry of liver tissue after I/R or I/R plus MTx. Oxidation of liver tissue was markedly increased in liver I/R control mice, while MTx significantly reduced 4-HNE lipid peroxidation at 2h of reperfusion (Fig. 2E-F). Sham-operated mice showed negligible amounts of 4-HNE lipid peroxidation.

To understand whether liver tissue was functionally improved in its bioenergetics and its regenerative capability, we measured fresh liver tissue ATP content and liver tissue hepatocyte growth factor (HGF). One hour of ischemia significantly decreased liver tissue ATP, as compared to sham-operated control mice (Fig. S1A). In response to liver reperfusion, ATP showed an initial small increase after 1h of reperfusion but decreased sharply by 2h of reperfusion (Fig. S1A). Injection of mitochondria in liver I/R, significantly increased liver tissue ATP content at 2h of reperfusion as compared to the I/R-only control group (Fig. S1A). We performed liver tissue ELISA for HGF (a key paracrine signalling cytokine in liver regeneration and mitogenesis) and found that liver I/R reduced tissue HGF, whereas injection of mitochondria in I/R restored tissue HGF to sham-operated control mice levels (Fig. S1B).

### Mitochondrial transplantation dampens systemic inflammation following liver I/R

Having demonstrated the beneficial effect of MTx on liver injury, we asked whether this translated into a salutary effect on systemic inflammation. We explored whether MTx might alter the circulating plasma cytokine profile, liver tissue cytokine expression and lung neutrophil infiltration following I/R (1h ischemia/2h of reperfusion). To confirm a reduction in systemic oxidative stress, we measured 8-isoprostane in mouse plasma, and found that MTx in I/R lowered this biomarker. Together, these data demonstrate that MTx is hepatoprotective in I/R, correlating with the reduction in hepatocellular oxidative stress (Fig. 3A). Consistent with others’ reports, liver I/R caused a significant rise in the pro-inflammatory cytokines, IL-6, and TNFα. As shown in Figures 3B and 3C, MTx significantly reduced pro-inflammatory IL-6 and TNFα responses (Fig. 3B-C). At the time point studied, liver mRNA expression of IL-6 and TNFα was not affected by MTx in I/R (Fig. 3D-E). By contrast, while MTx following I/R reduced systemic pro-inflammatory cytokines, circulating levels of the anti-inflammatory cytokine, IL-10, were significantly augmented following MTx in liver I/R (Fig. 3F). This was mirrored in the liver where tissue mRNA expression of IL-10 was significantly increased in mice that received MTx in liver I/R (Fig. 3G). Finally, plasma CCL2 (MCP-1), a robust chemoattractant for immune cells, which promotes hepatic injury in I/R via the CCR2 receptor, was significantly increased in I/R, and reduced in mice that received MTx (Fig. 3H).

**Fig. 3.**
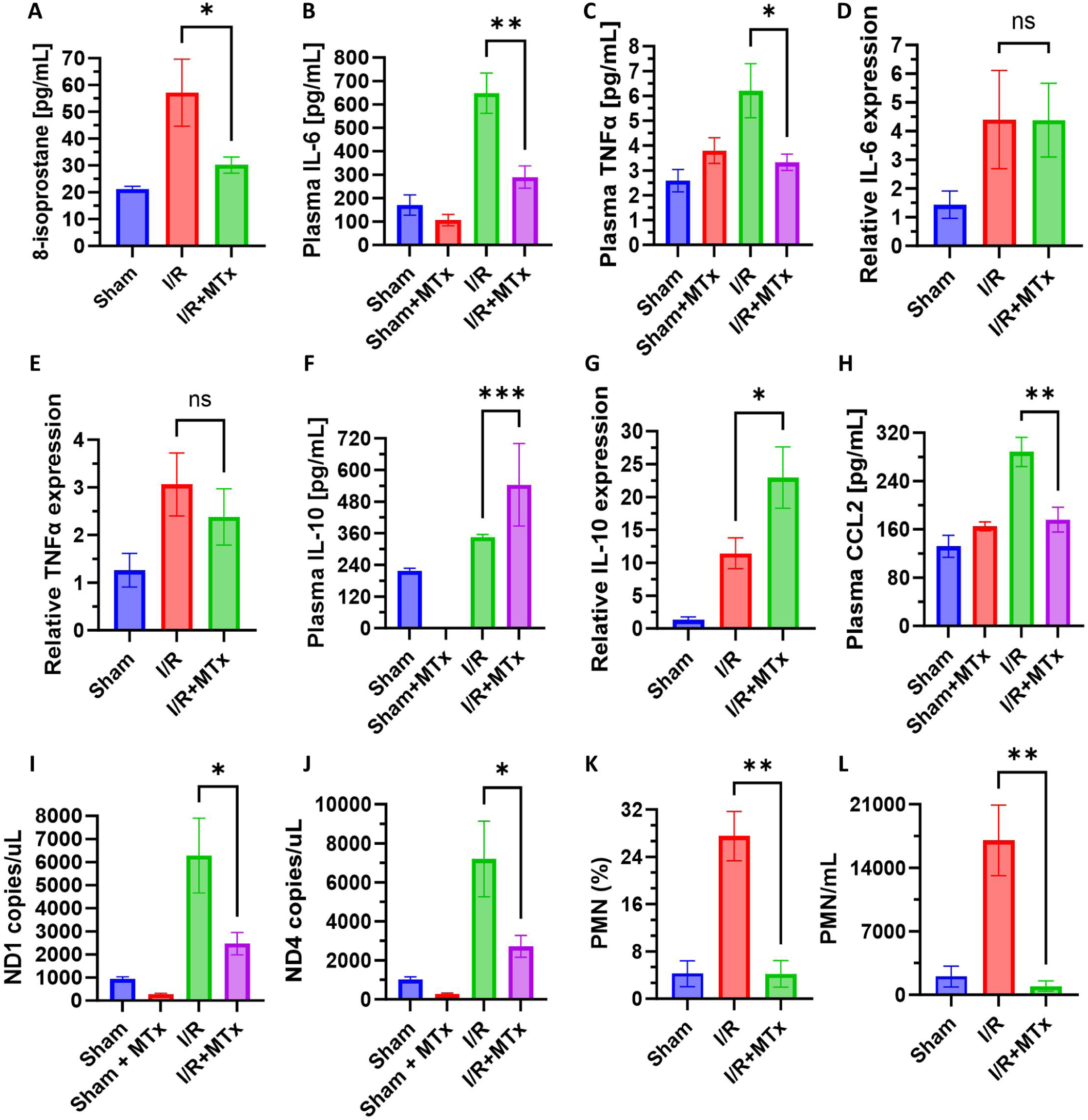
Mitochondrial transplantation dampens systemic inflammation following liver I/R. (A) MTx significantly lowered plasma 8-isoprostane, a marker of systemic oxidative stress, in liver I/R (p=0.002; n=5-8 mice per group). (**B-C**) MTx reduced circulating pro-inflammatory cytokines, plasma TNFα (p=0.009; n=3-6 mice per group) and IL-6 (p<0.0001; n=3-7 mice per group), in liver I/R. (**D-E**) Liver tissue expression of IL-6 and TNFα (n=7-10 mice per group). (**F**) MTx significantly lowered CCL2/MCP-1 in liver I/R (p=0.0002; n=3-8 mice per group). (**G- H**) Plasma IL-10 (p<0.0001; n=3-10 mice per group) and liver tissue IL-10 (p=0.0008; n=7-10 mice per group) expression showed that MTx significantly augmented this anti-inflammatory cytokine in liver I/R. (**I-J**) Circulating cell-free mitochondrial DNA (ccf-mtDNA), mt-ND1 (p=0.004; n=4-8 mice per group) and mt-ND4 (p=0.005; n=4-8 mice per group), were significantly increased by liver I/R, which were lessened in liver I/R + MTx mice. (**K-L**) Neutrophil exudation measured in lung BALF in liver I/R vs. liver I/R + MTx mice, showed lowered neutrophils in the alveolar space after treatment (p<0.01; n=3-6 mice per group).

Necrotic/injured cells release circulating cell-free mitochondrial DNA (ccf-mtDNA) which serve as a danger-associated molecular pattern (DAMP) and are pro-inflammatory. Liver I/R control mice showed a significantly increased mt-ND1 and mt-ND4 (genes encoded in the mitochondrial genome), as compared to sham-operated control mice (Fig. 3I-J), this suggested that there is a release of ccf-mtDNA into the systemic circulation following I/R injury. By contrast, injection of mitochondria in liver I/R significantly reduced these mito-DAMPs, as compared to liver I/R control mice (Fig. 3I-J).

Having shown a reduction in systemic inflammation by MTx, we asked whether distant organs might be affected. We studied lung inflammation by measuring neutrophil exudation into the alveolar space. As shown, the percentage and concentration of neutrophils in BALF was significantly reduced in mice injected with mitochondria in liver I/R, as compared to I/R-alone mice (Fig. 3K-L).

### Kupffer cells capture and acidify mitochondria in vivo

To study the fate of injected mitochondria, we performed liver intravital confocal microscopy (IVM) immediately after reperfusion and injection of mitochondria. To visualize cell populations in liver sinusoids, we labelled KCs (intravascular F4/80^+^), neutrophils (Ly6G^+^), and liver sinusoidal endothelial cells (CD31/PECAM-1^+^).

After 1h of liver ischemia, mice were prepared for IVM and mitochondrial injection was commenced. Imaging of the sinusoids after MTx in liver I/R revealed significant early localization of injected mitochondria (81±5s, Fig. 4A) on KCs, appearing within seconds of injection and persisting for the entire experimental period. Representative images are shown in Fig. 4B, and a video is presented in Movie S1. In the video, it is noteworthy that there is an initial bolus phenomenon (Movie S1) which diminishes over ∼15s, leaving transplanted mitochondria adherent to KCs over the duration of the experiment. As shown in Fig. 4C, time-dependent capture of mitochondria was quantitated. This revealed that mitochondria capture by I/R-subjected KCs was significantly higher throughout the experiment compared to sham-operated mice. This is clearly evident in the video files (Movie S1). Mitochondria adhered to KCs regardless of their functional status, as AntA-treated mitochondria were similarly captured (Fig. 4B). Representative videos of control mice are shown in Movie S2. Labelled mitochondria were not seen associated with endothelial cells or neutrophils (Fig. S2A), nor localized within the liver parenchyma (green autofluorescence channel for hepatocytes). In Fig. S2B, 3D z-stack images are shown of individual KCs with internalized aggregates of injected mitochondria and AntA-treated mitochondria.

**Fig. 4.**
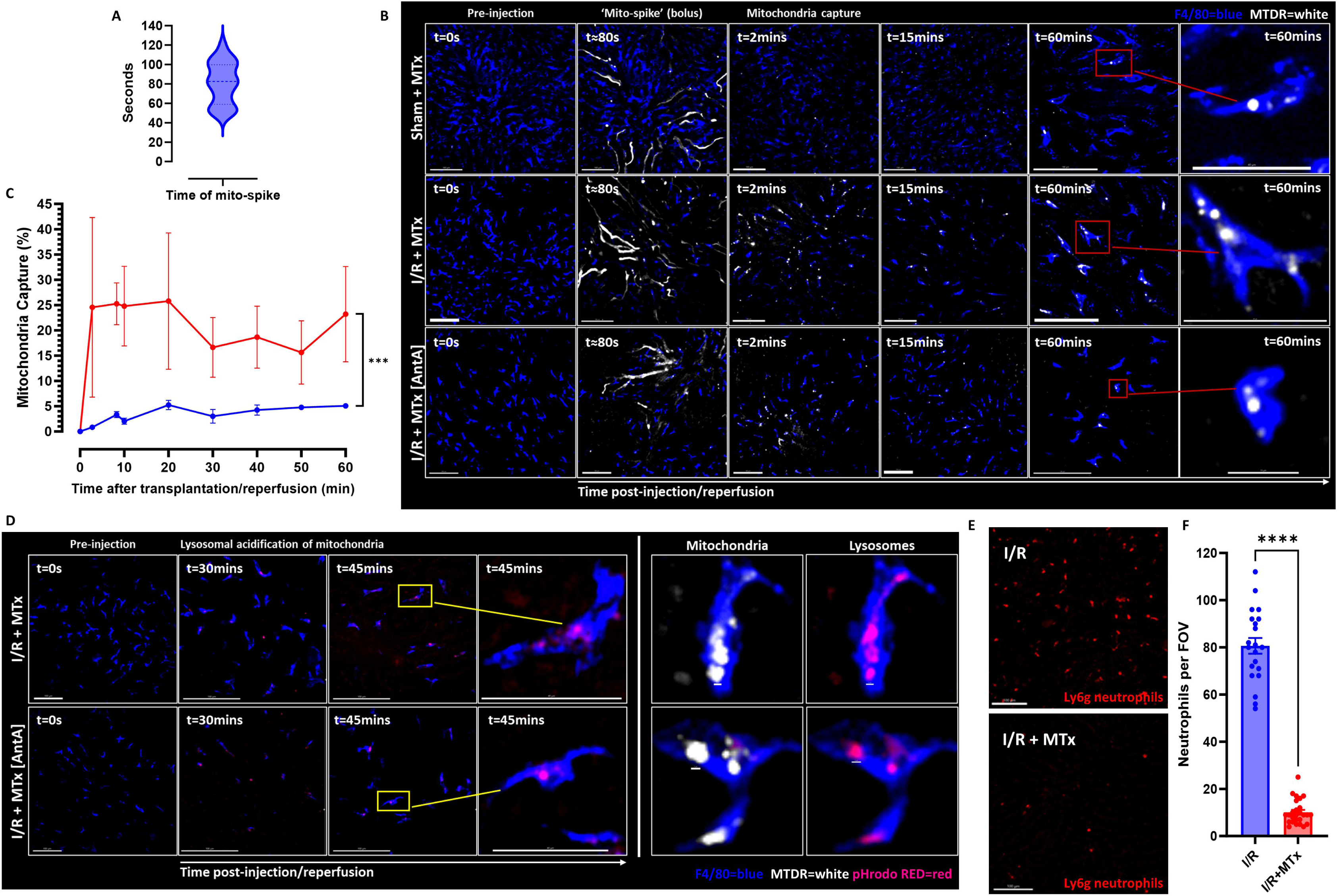
Kupffer cells capture and acidify mitochondria *in vivo*. (A) The transit time at which mitochondria reach the liver sinusoids after intrasplenic injection (i.e. the ‘mito-spike’ bolus). (B) Representative IVM images in C57BL/6J mice. F4/80^+^ KCs are in blue, and injected mitochondria are in white, with insets of single KCs. (**C**) Time-course quantification of mitochondria capture by KCs, showing significantly increased capture of mitochondria in I/R- subjected mice compared to sham-operated control mice (p<0.0001; n=3 sham mice, n=6 I/R mice. Movie S1 depicts mitochondria captured by KCs. (**D**) Representative IVM images in C57BL/6J mice. F4/80^+^ KCs are in blue, and captured mitochondria are in white, and acidified mitochondria are in red, with insets of single KCs showing acidified mitochondria in the lysosomal compartment. Movie S4 depicts KCs acidifying mitochondria *in vivo*. (**E-F**) Representative liver IVM images with LY6g^+^ neutrophils in red, revealing lowered neutrophils in mice that received MTx in I/R.

Lysosomal acidification of mitochondria by KCs, *in vivo*, was also studied. PHrodo is a succinimidyl ester dye which fluoresces red in the acidic environment of the lysosomal compartment and is commonly used for the detection of phagocytosis and endocytosis. We next double-labelled mitochondria with NHS ester AF647 and pHrodo RED. We found that both normal and AntA-treated mitochondria were acidified in lysosomal compartments by I/R- subjected KCs after mitochondrial injection (Fig. 4D). In Movie S3, we have shown representative acidification of mitochondria inside of KCs. Considered together, these studies show that KCs, subjected to I/R, quickly capture, internalize, and acidify mitochondria.

Finally, as a measure of local liver inflammation, LY6g^+^ neutrophils were quantified in IVM images. Neutrophils in the liver sinusoids were also lowered in mice injected with mitochondria in liver I/R, as compared to I/R-alone mice (Fig. 4E-F and Movie S4).

### Kupffer cells are required for the hepatoprotective effects of mitochondrial transplantation in liver I/R

Having demonstrated that KCs are the target cells for transplanted mitochondria, we conducted macrophage and KC depletion studies to determine whether these immune cells are necessary for mediating the hepatoprotective effects of MTx. First, clodronate liposomes were used to deplete macrophages in the liver (Fig. 5A). In our pilot studies, we found that the protocol of 1h ischemia followed by 2h reperfusion was lethal in macrophage-depleted animals, prompting us to modify the protocol to 0.5h ischemia/1h reperfusion. With this change in protocol, there was animal survival and maintenance of I/R-induced hepatocellular injury. As shown in Fig. 5B-C, injection of mitochondria in macrophage-depleted mice were not hepatoprotective and did not diminish the I/R-induced rise in proinflammatory cytokines/chemokines, including plasma TNFα, IL-6, and CCL2 (Fig. 5D-F).

**Fig. 5.**
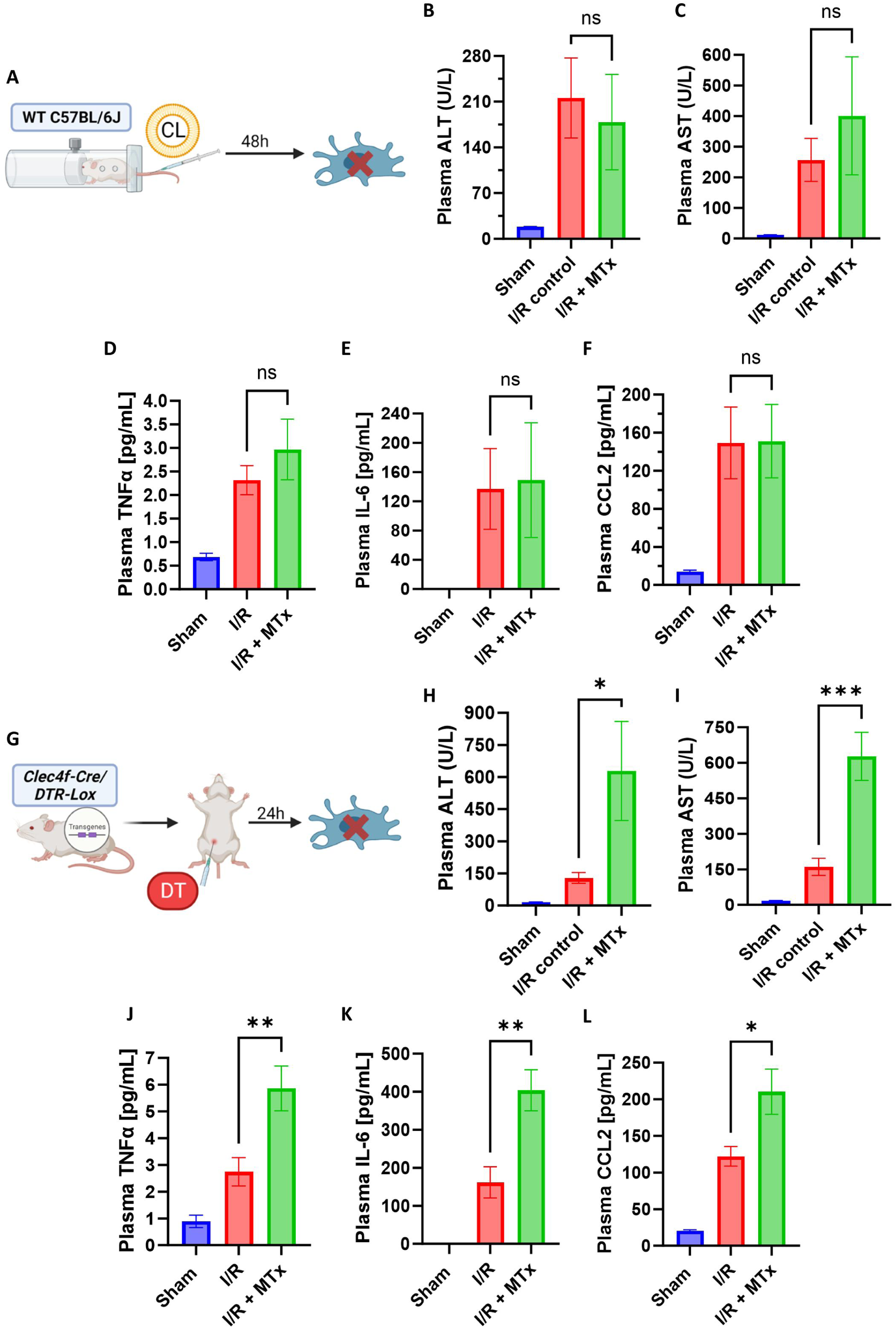
Kupffer cells are required for the hepatoprotective effects of mitochondrial transplantation in liver I/R. (**A-C**) Plasma ALT and AST in clodronate liposome, macrophage- depleted C57BL/6J mice, showing loss of the MTx hepatoprotective effects in liver I/R (0.5h ischemia/1h reperfusion) (p=0.06; n=3-5 mice per group). (**D-F**) Plasma pro-inflammatory cytokines/chemokines, TNFα (=0.0049; n=3-5 mice per group), IL-6 (ANOVA p=0.14; n=3-5 mice per group) and CCL2 (p=0.02; n=3-5 mice per group), in clodronate liposome, macrophage-depleted C57BL/6J mice, showing loss of the MTx anti-inflammatory effects in liver I/R. (**G-I**) Plasma ALT (p=0.014; n=5-7 mice per group) and AST (p<0.0001; n=5-7 mice per group) in diphtheria-toxin (DT), Kupffer cell-depleted Clec4f/iDTR mice, showing loss of the MTx hepatoprotection effects in liver I/R (0.5h ischemia/1h reperfusion) and increase in liver enzyme release. (**J-L**) Plasma pro-inflammatory cytokines/chemokines, TNFα (p=0.0002; n=5-7 mice per group), IL-6 (p<0.0001; n=5-7 mice per group) and CCL2 (p<0.0001; n=5-7 mice per group), in diphtheria-toxin (DT), Kupffer cell-depleted Clec4f/iDTR mice, showing an exacerbation of inflammatory responses.

In order to understand if KCs, *specifically*, have a role in mediating the hepatoprotection afforded by MTx in liver I/R, we utilized a diphtheria toxin (DT)-inducible, transient, depletion transgenic mouse model (Clec4f/iDTR mice) (Fig. 5G). Like clodronate-treated animals, animals with DT-mediated KC depletion had heightened susceptibility to I/R prompting us to shorten the protocol to that used for clodronate animals. In KC-depleted mice, liver I/R induced an increase in liver enzyme release, as compared to sham (DT-injected) mice. I/R-induced liver injury in these mice was not reduced by mitochondrial injection, but rather was exacerbated. (Fig. 5H-I). Similarly, in KC-depleted mice undergoing liver I/R, injection of mitochondria augmented plasma TNFα, IL-6 and CCL2 (Fig. 5J-L). Considered in aggregate, these macrophage and KC depletion studies revealed that KCs are required for the hepatoprotective effects of MTx in I/R.

#### Kupffer cells capture mitochondria through the CRIg/VSIG4 immunoreceptor and thereby attenuate liver I/R-induced injury and inflammation

Complement receptor of the immunoglobulin superfamily, CRIg (gene name *VSIG4*), is an immunoreceptor present on the plasma membrane of KCs. Since CRIg on KCs is required for the capture of gram-positive bacteria and other microbes,^20,39,40^ we hypothesized a potential role for CRIg in mitochondrial capture and resulting hepatoprotection.

In CRIg^-/-^ mice, there was a significant diminution in mitochondria capture by KCs in mice subjected to I/R compared to wild-type mice also subjected to I/R (Fig. 6A-B). Movie S5 shows impaired mitochondria capture in CRIg^-/-^ mice subjected to I/R. Importantly, CRIg^-/-^ mice undergoing liver I/R with MTx did not show hepatoprotection, as demonstrated by liver histology quantification (Fig. 6C-E) or by their ability to reduce plasma ALT and AST (Fig. 6F- G). Rather, injection of mitochondria in I/R-subjected CRIg^-/-^ mice aggravated liver injury (Fig. 6C-G). Additionally, in CRIg^-/-^ mice undergoing liver I/R with MTx, there was a significant rise in plasma TNFα, IL-6 and CCL2, compared to liver I/R mice (Fig. 6H-J). There were no significant differences in IL-10 in CRIg^-/-^ mice (Fig. 5K). Therefore, these data demonstrate that the absence of CRIg-dependent capture of mitochondria by Kupffer cells precluded the hepatoprotective and anti-inflammatory effects of MTx in liver I/R.

**Fig. 6.**
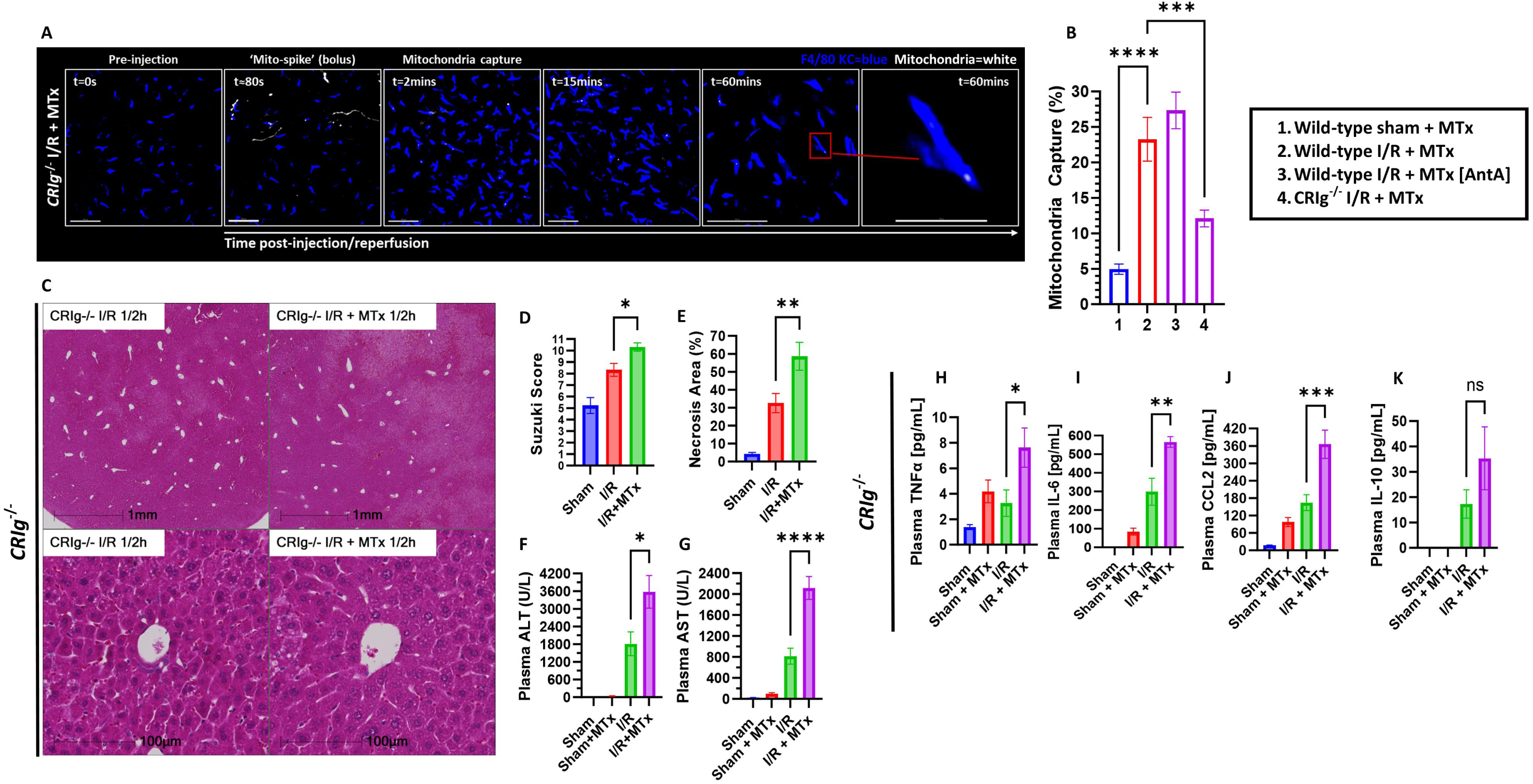
Kupffer cells capture mitochondria through the CRIg/VSIG4 immunoreceptor and attenuate liver I/R-induced injury and inflammation. (**A**) Representative IVM images in the mouse liver of CRIg^-/-^ mice subjected to liver I/R with MTx. F4/80^+^ KCs are in blue, and injected mitochondria are in white, with insets of single KCs. CRIg^-/-^ KCs have impaired mitochondria capture (n=6 mice). (**B**) Quantification of the percentage of KCs that captured mitochondria per FOV per mouse. Loss of CRIg results in significant diminishment of mitochondria capture by KCs (bar 4). (**C**) Representative liver histology in CRIg^-/-^ mice subjected to liver I/R ± MTx, showing no hepatoprotection. (**D**) Suzuki liver injury score quantified by 3 blinded observers per CRIg^-/-^ mouse. CRIg^-/-^ mice undergoing liver I/R with MTx showed no hepatocellular protection and increased injury (p<0.0001; n=5-7 mice per group). (**E**) Quantification of necrosis area percentage in CRIg^-/-^ mice. CRIg^-/-^ mice undergoing liver I/R with MTx showed higher necrosis area, compared to liver I/R mice (p<0.0001; n=5-7 mice per group). (**F-G**) Plasma ALT and AST in CRIg^-/-^ mice, showing loss of the MTx hepatoprotection in liver I/R, and depicting an exacerbation of liver enzyme release (p<0.0001; n=6-7 mice per group). (**H-J**) Plasma cytokines/chemokines in CRIg^-/-^ mice. Loss of CRIg resulted in the loss of the anti-inflammatory effects of MTx in liver I/R, as shown by plasma TNFα (p=0.005; n=3-7 mice per group), IL-6 (p<0.0001; n=6-7 mice per group) and CCL2 (p<0.0001; n=3-7 mice per group), rather, there is an increase in the inflammatory response with MTx. (**K**) In CRIg^-/-^ mice, plasma IL-10 showed no significant differences between groups (p=0.01; n=3-7 mice per group).

## DISCUSSION

I/R underlies the pathophysiology of several clinical settings including stroke, myocardial infarction and organ transplantation. I/R is characterized by the development of mitochondrial dysfunction which leads to both a defect in cellular bioenergetics but also is profoundly pro-inflammatory, in part through the elaboration of mitochondrial reactive oxygen species. Strategies aimed at restoring mitochondrial homeostasis represent a potential therapeutic approach. MTx has been proposed as an opportunity to achieve this. In the current studies, we demonstrate that mitochondria isolated from skeletal muscle and delivered to the liver via the portal vein at the initiation of the reperfusion phase are able lessen hepatocellular inflammation and injury. *In vivo* microscopic imaging provides novel insight into the mechanism by definitively demonstrating that injected viable mitochondria are captured and internalized by KCs in a CRIg-dependent fashion. The requirement for this interaction between mitochondria and KCs was demonstrated by the loss of hepatoprotection when these cells were depleted.

Two prior studies have reported a hepatoprotective effect of MTx in liver I/R,^8,9^ but have been observational in nature and have not specifically identified the KC as the critical cell contributing to protection. Portal vein delivery of melatonin-treated isolated mitochondria has been shown to be protective in a rat liver I/R model,^8^ but detailed studies of localization were not performed. Glaumann *et al*. (1975), and Martin and colleagues investigated the uptake and lysosomal digestion of intravenously injected isotope-labelled mitochondria by rat KCs, as a means of studying the disposition of injected mitochondria, rather than exploring their potential hepatoprotective effects.^41–43^ Our studies demonstrate a clear linkage between KC capture of transplanted mitochondria and the resultant hepatoprotection. A number of mechanisms have been proposed for the cytoprotective effects of MTx, most notable is the internalization of mitochondria by the target cell, e.g. the cardiomyocyte, via actin-dependent endocytosis with consequent beneficial effects in the recipient cell mediated by escape of the mitochondria from the endolysomal compartment and fusion with endogenous mitochondria.^44–46^ This results in improved bioenergetics, mitochondrial function, and respiration.^44–46^ The finding that live mitochondria but not AntA-treated mitochondria were able to exert protection suggest that a bioenergetic effect on the KC might be important. Rather, we have focused on the downstream impact of the KC-mitochondrial interaction. As we show, interaction of mitochondria with KCs causes a shift in the cytokine milieu from one which is proinflammatory to one which is anti-inflammatory and influences the progression of inflammation and injury in nearby cells through paracrine signalling in the liver microenvironment.

A number of potential mechanisms may have contributed to the shift in KCs from pro-inflammatory to anti-inflammatory following interaction with transplanted mitochondria. One might relate to an immunophenotypic shift from M1-like KC phenotype induced by I/R to an anti-inflammatory M2 KC phenotype. In our studies, the CRIg receptor on the surface of KCs was shown to be required for uptake of mitochondria and induction of the hepatoprotection. Generally, CRIg-mediated signalling is anti-inflammatory, and inhibits both pro-inflammatory macrophage and T-cell activation.^20,47–50^ At the molecular level, CRIg blocks TLR4/NF-κB- mediated inflammation signalling, and activates PI3K/AKT/JAK2/STAT3 anti-inflammatory programs, thereby inhibiting M1 macrophage polarization and promoting M2 polarization towards M2, and represses NLRP3 inflammation.^47,51–54^ One alternative explanation is that MTx might transition endogenous mitochondrial respiration from glycolysis to oxidative phosphorylation.^55,56^ This metabolic shift has been shown to be associated with pro- and anti-inflammatory macrophage phenotypes respectively,^57,58^ and hence might account for the hepatoprotective effect of MTx. The uptake of mitochondria by rat myoblasts and mouse fibroblasts has also been shown to improve oxidative phosphorylation and reduce glycolysis.^56^ Further, the treatment of bone marrow-derived macrophages with mitochondria lowered M1 polarization and release of pro-inflammatory cytokines,^55^ similar immunophenotypic switching has been reported in microglia.^59^ Finally, it is possible that KCs sense and process active and inactive mitochondria differentially, similar to how macrophages sense microbial ‘viability- PAMPs’ and tailor the inflammatory response.^60,61^ While the focus of this manuscript has been on KC function, the magnitude of the inflammatory response and hepatocellular necrosis after liver injury, is augmented by the infiltration of pro-inflammatory Ly-6C^high^ circulating monocyte-derived macrophages (MoMFs) attracted by the release of CCR2 chemokines from KCs.^62,63^ Reduced CCL2 from KCs may also have lessened cellular infiltration and injury. Clearly, further studies are required to better understand the change in KC function following capture and internalization of exogenous mitochondria.

MTx was performed at the end of the ischemic phase, at the onset of reperfusion. This timing has relevance to the clinical setting where the onset of the reperfusion phase is more predictable. In our studies, mitochondria were isolated from hindlimb skeletal muscle of a separate animal in a process taking approximately 3h. This may be suitable for some settings, e.g. transplantation, where timing can be controlled but not for others, e.g. traumatic hemorrhagic shock, where the scenario is unpredictable. Ideally, having mitochondria available for use, similar to a blood bank, would be optimal. For example, Cloer and colleagues used frozen stored allogeneic mitochondria in both porcine and human *ex vivo* lung perfusion and showed that MTx reduced proinflammatory cytokine secretion and improved pulmonary vascular resistance.^64^ Using a protocol of storing mitochondria in trehalose-containing solution at -80°C, function was retained for one year.^64^

In summary, our studies demonstrate a novel mechanism whereby MTx exerts hepatoprotection in a model of I/R injury. In addition to future studies aimed at further elucidating key signaling pathways, a focus on streamlining the process of mitochondrial isolation and preserving mitochondrial function following recovery may position this as a potential therapeutic intervention in a range of disease processes whose common pathogenesis is I/R. Finally, translation of these findings to large animal models would position MTx as a novel therapeutic in surgery.

## Funding

ANM was supported by funding from the Department of Surgery, St. Michael’s Hospital; Mitochondria Innovation Initiative (Mito2i), University of Toronto; Medicine by Design, University of Toronto; Institute of Biomedical Engineering, Science and Technology (iBEST), Unity Health Toronto; Faculty of Science, Toronto Metropolitan University; Institute of Medical Science, University of Toronto; and the Shock Society. This work was funded through research grants awarded to ODR by Canadian Institutes for Health Research (CIHR, #23766), and Defence Research & Development Canada (W7719-225559 and W7714-145967). HN was awarded a CIHR grant (#389035). DK was supported by the Vanier Canada Graduate Scholarship by CIHR.

## Acknowledgments

The authors thank Drs. Pamela Plant, Caterina Di Ciano-Oliveira, Monika Lodyga, Xiaofeng Lu, Dario Bogojevic from the Research Core Facilities at the Keenan Research Centre for Biomedical Science, Unity Health Toronto, for their methodological expertise and support services. The authors thank Dr. Lindsey Fiddes and Yan Chen at the Microscopy Imaging Laboratory, Temerty Faculty of Medicine, University of Toronto, for their methodological expertise in transmission electron microscopy. The authors thank Donna Lyons, Danielle Bince, Lauren Sturrock, Kaili Metsala and Amanda Hillier from the Research Vivarium at the Keenan Research Centre for Biomedical Science, Unity Health Toronto, for their diligent animal care, colony maintenance, and breeding of mice.

## Author contributions

Author contributions are listed according to the CRediT model: Conceptualization: ODR, ANM, MA, AK, PK Methodology: ANM, BAD, JL, CMV, DK, MA, RG, WB, AG, RSW, KJ, RP, MJ Formal analysis: ANM, DK, MA Data curation: ANM Validation: ANM, BAD, JL, CMV, DK, MA Investigation: ANM, BAD, MA, PK, ODR Visualization: ANM Funding acquisition: ODR Project administration: ODR Software: ANM, DK, MA, RG, WB, AG, RSW, KJ, RP Supervision: ODR Resources: ODR, PK, HZ, AK, HN Writing – original draft: ANM, ODR Writing – review & editing: all authors

## Competing interests

The authors have no disclosures or COI of relevance to this publication.

## Video captions

**Movie S1**. **Capture of transplanted mitochondria by I/R-subjected Kupffer cells, *in vivo*.** Representative IVM videos of C57BL/6J wild-type mice. Groups include Sham+MTx and liver I/R+MTx and liver I/R+MTx [AntA]. 1h of ischemia, followed by liver reperfusion and immediate intrasplenic injection of mitochondria (white), Kupffer cells (blue).

**Movie S2**. **Mitochondria staining and injection controls for IVM studies**. Representative IVM videos of C57BL/6J wild-type control mice: unlabeled mitochondria and vehicle. F4/80^+^ Kupffer cells are shown in blue.

**Movie S3**. Acidification of mitochondria by I/R-subjected Kupffer cells, *in vivo*. PHrodo^+^ mitochondria (red) and F4/80^+^ Kupffer cells (blue).

**Movie S4**. **Neutrophils in the mouse liver sinusoids following MTx in liver I/R**. Video 1 is a liver I/R mouse and video 2 is a liver I/R + MTx mouse. LY6g^+^ neutrophils are shown in red.

**Movie S5**. **Impairment of mitochondria capture in CRIg^-/-^ mice subjected to liver I/R**. Transplanted mitochondria are white and Kupffer cells are blue.

## Notes

### Competing Interest Statement

The authors have declared no competing interest.

