## Supplementary methods, supplementary figure legends, supplementary tables for "Mitochondrial transplantation: A novel therapy for liver ischemia/reperfusion injury"

**Mukkala *et al*. 2024.**

**SUPPLEMENTARY INFORMATION**

**Supplementary Methods**

**Intravital confocal microscopy and quantification**

Intravital microscopy (IVM) was customised from Peiseler and David *et al*., Marques *et al.* and Surewaard *et al.*^1-3^ Mice received cell-labelling antibodies retro-orbitally 30mins prior to liver ischemia. CD31/PECAM-1 for liver sinusoidal endothelial cells (LSECs), and Ly6G for neutrophils. Kupffer cells, were labelled with F4/80 and as confirmatory control staining with TIM4 or CRIg/VSIG4 (Table S1), and hepatocytes on the green autofluorescence channel. C57BL/6J and CRIg^-/-^ mice were subjected to 1h of liver ischemia, as described. Intrasplenic injection of mitochondria was conducted immediately after clamp removal/reperfusion of liver. Mitochondria were stained with 100nM of MTDR, and washed twice with RB at 9000xg for 10mins at 4°C. As negative controls, vehicle RB and unlabelled mitochondria were injected. As a positive control for phagocytosis studies, methanol-killed *S. aureus* BioParticles-conjugated to pH-RODO RED were acquired (Table S1) and labelled as described by Surewaard *et al.*^3^

To prepare the mouse liver for IVM, part of the abdominal wall and peritoneum were removed with high-temperature cautery (Table S1). The mouse was placed in a lateral position on the heating stage to maintain body temperature at 37°C. Then, the left lobe of the liver was exteriorized and placed onto a customized glass coverslip window, using fine saline-wet cotton swabs, using a no-touch technique, the liver was covered with sterile saline soaked Kimwipes in order to decrease movement due to mouse breathing. All other abdominal organs were covered with sterile-saline cotton gauze to prevent dehydration. Mice received constant inhaled anesthetic by a mobile isoflurane-oxygen gas mixture unit.

Specific details of the confocal microscope setup can be found at Peiseler and David *et al*.^2^ Briefly, acquisition of images was on an inverted spinning-disk confocal microscope (IX81; Olympus). The confocal light path (WaveFx; Quorum Technologies) had a modified CSU-10 head (Yokogawa Electric Corporation). Laser excitation wavelengths were 405nm, 488nm, 561nm, 642nm, 735nm (Cobolt, Stockholm, Sweden). A 512x512 pixel back-thinned EMCCD camera was utilized for detection (C9100-13, Hamamatsu, Bridgewater, NJ). Volocity software (Quorum) was used to drive the microscope and for acquisition. Time-lapse videos were recorded from the time of reperfusion/injection (t=0h) until sacrifice. Videos were recorded at 10X or 20X magnification, ~5.5mins per FOV, and 8-12 FOVs per mouse. Z-stacks were achieved after mice were sacrificed by ketamine/xylazine overdose. Image files and time-lapse videos were exported from VOLOCITY for analysis in IMARIS.

Each mvd2 image/video file was converted to TIFF using Fiji/ImageJ. Images and videos were opened in IMARIS (v9.8.2), and thresholding was applied based on control mice. Kupffer cells, in liver sinusoids, were identified as being F4/80^+^ (which were also TIM4^+^ and CRIg^+^), injected and captured mitochondria were identified as being MTDR^+^ and acidified mitochondria were identified as being pH-RODO RED^+^. The IMARIS cell and spot counting functions were utilized to count the number of Kupffer cells (>20um), mitochondria (<4um), neutrophils (<20um) for each second of each FOV in each mouse. Captured mitochondria were defined as being arrested by Kupffer cells for 2.5mins. The number of captured mitochondria and the number of Kupffer cells with captured mitochondria were counted for each FOV in each mouse. The percentage of Kupffer cells that had captured mitochondria was calculated per FOV – this was defined as mitochondria capture. Video FOVs that contained excess movement (due to breathing), or background signal were excluded. A total of 325 FOVs were analyzed, which contained ~2000mins, and spatially represented ~6.26x10^7^ voxels.

**Quantitative reverse transcription polymerase chain reaction (qPCR)**

Liver tissue was excised from mice and immediately placed in ice cold 0.9% sterile saline and washed thrice. The liver tissue was cut into 3-5mm^3^ and snap frozen in liquid nitrogen and stored at -80°C. RNA was extracted using the Qiagen RNeasy Mini Kit. 20uL of RLT buffer (+BME) was added per mg of liver tissue. Briefly, 35mg of liver tissue was weighed and homogenized in sterile Eppendorf mortar and pestle. The tissue lysate was passed through a sterile 26-gauge needle and through the Qiashredder at 21,100 x g for 2min at room temperature. One volume of 50% EtOH was added to the supernatant and mixed by pipetting. The rest of the RNA extraction was according to the Qiagen RNeasy Mini Kit with a final elution volume of 40uL. RNA samples were diluted 15-fold in qPCR-grade ddH_2_O, and RNA concentration was quantified using the Qubit spectrophotometer (ThermoFisher). 1716±652 ng/uL of RNA was extracted. RNA samples were assessed for quality using the Agilent 2100 Bioanalyzer. Only samples with RIN (RNA integrity number) ≥8.0 were used for qPCR assays and the average RIN was 9.0±0.3. One microgram of RNA was DNase-treated using a DNase I kit (Table S1). cDNA was synthesized using iScript cDNA synthesis kit (Table S1). Each primer set was validated by generating amplification efficiency curves, melting curves and DNA product was run on 1.5% agarose gel to confirm amplicon size. qPCR reactions were in 10uL total volume. The reaction was as follows: 5uL PowerSYBR GREEN PCR master mix (Table S1), 300nM forward primer, 300nM reverse primer, 2.4uL qPCR-grade ddH_2_O and 2uL of cDNA template. The thermocycler settings were as follows on the QuantStudio 7 system (Thermo Fisher Scientific): 50°C (2mins), 95°C (10mins), 40 annealing cycles between 95°C (15s) and 60°C (1min) then 4°C, with all ramps at 1.6°C/s. Data was exported and analyzed by 2^-∆∆Ct^ Livak method. Data was normalized to sham mice. Primers are listed in table S2.

**Droplet digital polymerase chain reaction (ddPCR) of circulating cell-free mtDNA**

ddPCR was adapted from previously established methods.^4^ Mouse blood was centrifuged at 2500 x g for 10mins at 4°C, and the plasma supernatant was collected and stored at -80°C. DNA was extracted from 20uL of mouse plasma using the Qiagen QIAamp DNA Micro Kit (with carrier RNA). DNA concentration was quantified using the NanoDrop 2000 and 35±5ng/uL of ccf-DNA was present. The ddPCR assay was validated using thermal gradient, ccf-DNA and primer dilution series. 1/100 ccf-DNA dilution was used in ddPCR assays. The ddPCR reaction consisted of 10uL EvaGreen SuperMix, 250nM forward primer, 250nM reverse primer, 6.7uL of ultrapure ddH_2_O and 2.2uL of 1/100 diluted ccf-DNA template. Primer annealing temperature was determined to be 60°C. Target genes for this assay were mt-ND1 and mt-ND4, and housekeeping genes were B2M and B-actin. Master mixes were prepared for each target gene and thoroughly vortexed at maximum speed for 15s. The reaction was loaded in duplicate as follows: 19.8uL master mix plus 2.2uL 1/100 diluted ccf-DNA template. Plates were heat-sealed with aluminum covers and again thoroughly vortexed for 1min. Plates were centrifuged at 1000 x g for 2mins, and droplets were generated using the automated droplet generator (Bio-Rad, QX200 AutoDG). Plates underwent PCR in the C100 BioRad thermocycler using the following settings: 95°C denaturation (5min), 40 annealing cycles between 95°C (30s) and 60°C (1min), 4°C (5mins), 90°C (5mins), and 4°C (5mins). Droplet nanoreactions were read using the BioRad droplet reader (Bio-Rad, QX200 Droplet Reader). Data was exported for analysis as absolute gene copies per uL. Primers are listed in table S2.

**Transmission electron microscopy of isolated mouse skeletal muscle mitochondria**

TEM was described previously.^5^ Mitochondria were pelleted at 9000 x g for 10mins at 4°C. 1mL of primary fixative (4% Paraformaldehyde plus 1% Glutaraldehyde in 0.1 M phosphate buffer, pH = 7.2) was added to the pellet without disruption for at least 24h at 4°C. Mitochondria were stained in 0.15% tannic acid in 0.1M pH 7.2 PB for 20min for contrast enhancement and washed three times in 0.1M pH 7.2 PB, 20mins per wash to remove unfixed aldehydes. Mitochondria were then postfixed in secondary fixative (1% Osmium Tetraoxide buffered with 0.1 M Phosphate, pH = 7.2) for 1h at room temperature and washed three times in 0.1M pH 7.2 PB, 20mins per wash. Mitochondria were dehydrated in an ethanol series over 3h as follows: 30% 2 changes with 10min each, 50% 2 changes with 10min each, 70% 2 changes with 10min each, 90% 2 changes with 10min each, 100% 3 changes with 15min each, and propylene oxide 2 changes with 15min each. Mitochondria were infiltrated with Epon resin as follows: 1-part Epon resin mixed with 2 parts propylene oxide for 1h using an agitator; 2 parts Epon resin mixed with 1 part propylene oxide for 3h using an agitator; 100% Epon overnight using an agitator; one change with fresh resin agitated for 2h. Mitochondria, in Epon resin, were then polymerized at 60°C for 48h. Ultrathin 80-100nm sections were cut. Samples were examined utilizing the Talos L120C transmission electron microscope (TEM) system, and 5 random images per sample were captured with the high resolution 4 K × 4 K CETA CMOS camera (ThermoFischer Scientific). TEM was conducted at the Microscopy Imaging Laboratory at the University of Toronto.

**Mitochondria labelling and nanocytometry**

Mitochondria were labelled with MitoTracker DEEP RED (MTDR) FM, or JC-1, or pH-RODO, ATPB (Complex V, F_1_F_0_ ATP synthase) AlexaFluor 647, NHS ester AlexaFluor 647 (Table S1). Mitochondria were incubated with gentle end-over-end rotation or gentle rocking at 4°C for 30mins. Mitochondria were washed twice in respiration buffer by centrifugation at 9000 x g for 10mins at 4°C and resuspended in respiration buffer for mitochondrial transplantation studies. For intravital microscopy acidification studies, mitochondria were double-labelled with NHS ester AlexaFluor 647 and pH-RODO RED and fluorescence was confirmed by suspending the mitochondria in a pH=4 versus pH=7 respiration buffer then performing nanocytometry or mitochondria transplant in I/R for intravital microscopy, as described. A titration was done for MitoTracker DEEP RED FM and pH-RODO RED to optimize fluorescence signal on intravital microscopy (100-400nM MTDR and 25-25000ng/mL pH-RODO). Single stain, and unlabelled controls were conducted.

Mitochondrial nanocytometry was adapted from MacDonald *et al*.^6^ Labelled or unstained mitochondria were suspended in respiration buffer. The SONY SP6800 flow cytometer was size calibrated using SpheroTech fluorescent beads for detection of mitochondria (Table S1). Size gating was performed with the lower diameter limit at 700-900nm and an upper diameter limit at 1700-2200nm. NILE RED fluorescent beads (1700-2200nm, mean=2100nm) and JADE GREEN fluorescent beads (700-900nm, mean=830nm) were gated, followed by a mitochondria size gate which encompassed both of these diameter ranges. 2.5x10^6^ bead events were collected in order to closely define the nano-scale mitochondria size gate by forward and side scatter (area). This range was deemed sufficient and appropriate given that mitochondria consistently fall within the <4um range on the Coulter particle counter with an average size of 1160±14.35nm and 938±170nm on the DLS ZetaSizer. The mitochondria population was gated as being MTDR+ (100nM) as compared to unstained mitochondria control. Mitochondria were gated for analyses of membrane potential with JC-1. Controls were conducted with vehicle respiration buffer, unstained mitochondria, LN_2_-disrupted, or AntA-treated (50uM). Data was exported and analyzed on FlowJo (v10).

**Liver tissue ATP**

Extraction of ATP from fresh mammalian tissue was modified from Chida *et al. ^7^* Briefly, liver tissue was excised and washed three times in ice-cold PBS. 160mg of liver tissue was weighed. Liver tissue was thoroughly homogenized in 20x volume of ice-cold phenol-saturated TE (Invitrogen) using a glass 3mL Dounce homogenizer. The homogenate was mixed by pipetting and 1mL was aliquoted into an Eppendorf tube on ice. 200uL of pure chloroform and 150uL of distilled water were added. The mixture was vortexed for exactly 30s at maximum speed then centrifuged at 20,000 x g for 10mins at 4°C. Three layers form after this centrifugation, only the topmost supernatant was collected and placed on ice. The ATP extract was diluted 1000-fold in ultraclean water and ATP was measured immediately. ATP concentration was quantified using a luciferase bioluminescence assay kit (Table S1) in an opaque white-bottom 96-well plate. The plate was read at 560nm RLU at 28°C in a luminometer (M5e, Molecular Devices). Data were calculated to the ATP standard curve provided in the kit.

**Lung bronchoalveolar lavage fluid (BALF) neutrophil quantification**

BALF was collected by inserting a catheter in the trachea of terminally anesthetized mice. The left lung was lavaged 3 times with 500uL sterile PBS. The lavage fluid was centrifuged at 1000xg for 5 min at 4°C, and the supernatant collected and stored in -80°C. The cell pellet was used for total cell count and Cytospin analysis (400xg for 5mins onto slides) followed by H&E staining (neutrophils and macrophages). Using the Zeiss Axio Scan (v2.1) system, images of entire slides were acquired. Image files were exported for blinded analysis in ZEN (Blue 2012). Percentage neutrophils in BALF and PMN per mL was calculated.

**Western blot of isolated mitochondria**

Western blotting was conducted as previously described.^8^ Mitochondria were lysed in lysis buffer and briefly mixed, then incubated on ice for 20mins. After centrifugation at 12,000 RPM (13,800 x g) for 10min at 4°C, the supernatant was collected and aliquoted for storage at -80°C until use. Protein concentrations were determined by Bio-Rad DC Protein Assay (Bio-Rad, Hercules, CA). Western blotting was conducted as described above for liver tissue, but with 20ug of protein per lane. Lysis buffer: 10 mM NaCl, 30 mM HEPES, 20 mM NaF, 1 mM EGTA, 1% Triton X, 1 mM sodium orthovanadate, and complete protease inhibitor cocktail (Roche Diagnostics, Mannheim, Germany). 10-40ug of protein sample was loaded onto 12% sodium dodecyl sulfate–polyacrylamide gel for electrophoresis (SDS-PAGE), and electrophoretic transfer was performed onto nitrocellulose membranes (Bio-Rad). Membranes were blocked on a rocking platform for 1h at room temperature in 5% skim milk in tris-buffered saline Tween 20 (TBST). Membranes were probed with primary antibodies (antibodies/dilutions listed in Table S1) diluted in 5% skim milk-TBST and incubated overnight at 4°C on a rocking platform. After 3 washes in TBST, membranes were incubated with appropriate secondary horseradish peroxidase antibodies (Table S1) for 1 h at room temperature. After 3 TBST washes, membranes underwent enhanced chemiluminescence (Ultrascence ECL Western Blot Substrate, Frogga Bio) followed by exposure in the ChemiDoc Touch Imaging System (Bio-Rad).

**Hepatic and splenic macrophage depletion flow cytometry**

Flow cytometry was described previously,^9^ and adapted for our purposes as follows. Naïve wild-type or transgenic mice were sacrificed by carotid artery exsanguination and thoroughly bled. The gallbladder was removed, and the liver and spleen were excised and placed into 10mL of HBSS at room temperature. The livers were homogenized through a 100μm nylon mesh filter to create a single-cell liver suspension. The homogenate was centrifuged twice at 50 x g, 3mins each at RT, and each time the hepatocyte pellet was discarded. The liver non-parenchymal cell (NPC) fraction was pelleted at 600 x g for 6mins. The supernatant was discarded and the LNPC pellet was resuspended in PBS. Live/dead cell control was conducted by heat-killing an aliquot of LNPCs for 10mins at 65°C and then placed on ice for 1min. Then, 50uL of all samples were stained with 100uL of 1/250 Zombie aqua for 15mins at RT, protected from light. The samples were washed with FACS Buffer and fixed in 4% paraformaldehyde (PFA) for 15mins at RT, protected from light. Then, all samples were washed twice in FACS buffer and Fc receptors were blocked with TruStain FcX (Table S1) Antibody. Cell samples were stained at 1ug/mL with F4/80 BV711 (Table S1) for 1h at RT, protected from light, then washed with FACS buffer. Cell samples were washed with permeabilization buffer. Cell samples were then stained with CD68 APC/Cy7 (Table S1) at 0.5ug/mL in permeabilization buffer for 1h at RT, protected from light. Cell samples were washed in permeabilization buffer and washed in FACS buffer. Cells were finally resuspended in FACS buffer. All samples were acquired on the SONY SP6800 flow cytometer. Kupffer cells and splenic resident macrophages (SRM) were identified as being double positive for CD68 and F4/80.

**SUPPLEMENTARY FIGURE LEGENDS**

**Fig. S1. Mitochondrial transplantation improves liver tissue ATP and HGF following I/R.** (**A**) Fresh liver tissue ATP in C57BL/6J mice subjected to liver I/R with or without MTx. Liver ischemia perturbs ATP, and at t=2h after reperfusion, MTx in liver I/R improves ATP content, compared to the liver I/R control group (p<0.0001; n=3-9 mice per group). (**B**) Liver tissue HGF, a biomarker of regeneration/mitogenesis, was restored by MTx in liver I/R, whereas liver I/R control mice showed a reduction in HGF (p<0.0001; n=6-12 mice per group).

**Fig. S2. Transplanted mitochondria localize to I/R-subjected Kupffer cells in the liver sinusoids and are internalized.** (**A**) Representative intravital confocal microscopy images of PECAM/CD31+ liver sinusoidal endothelial cells (LSEC), Ly6g^+^ neutrophils and F4/80^+^ Kupffer cells, and transplanted mitochondria (MTDR^+^). Mitochondria localize to Kupffer cells, and not to other cell types. (**B**) 3D z-stack images showing single KCs with internalized mitochondria. F4/80^+^ KCs are in blue, and MTDR^+^ mitochondria aggregates are in white.

**Table S1. Key resources, materials, mice, and reagents**

| **REAGENT/RESOURCE** | **SOURCE** | **IDENTIFIER** | **Dilution/dose** | **Purpose(s)** |
| --- | --- | --- | --- | --- |
| **Mice** |  |  |  |  |
| C57BL/6J mice (wild-type) | Jackson Laboratories | 000664 | N/A | Animal studies |
| C57BL/6J-Clec4f^em1(cre)Glass/J^ (Clec4f-Cre-tdTomato) | Dr. Heyu Ni Lab (originally from JAX) | 033296 | N/A | Animal studies |
| C57BL/6-Gt(ROSA)26Sor^tm1(HBEGF)Awai/J^ (ROSA26iDTR) | Dr. Heyu Ni Lab (originally from JAX) | 007900 | N/A | Animal studies |
| CRIg Knockout Mice (CRIg^-/-^) | Genentech | N/A | N/A | Animal studies |
| **Antibodies** |  |  |  |  |
| Anti-mouse VSIG4 (CRIg) AlexaFluor488 | eBioscience, Invitrogen, Thermo Fisher Scientific | Clone NLA14  Cat#53-575-282 | 1/100  5ug/mL | IF and IVM |
| Anti-mouse F4/80-BV711 | Biolegend | Clone BM8  Cat#123147 | 1ug/mL | FC |
| Anti-mouse CD68 APC/Cy7 | Biolegend | Clone FA-11  Cat#137024 | 0.5ug/mL | FC |
| Anti-F_1_F_0_-α mouse mAb | Calbiochem | AP1036 | 1/1000 | WB |
| TOM20 | Cell Signalling | 42406S | 1/2000 | WB |
| CRIg/VSIG4 | Abcam | ab252933 | 1/1000 | WB |
| GAPDH | Cell Signalling | 5174S | 1/40,000 | WB |
| AffiniPure donkey anti-rabbit IgG HRP | Jackson Immuno-Research | #711-035-152 | 1/1000 – 1/40,000 | WB |
| AffiniPure goat anti-mouse  IgG light chain HRP | Jackson Immuno-Research | #115-035-174 | 1/1000 – 1/40,000 | WB |
| AffiniPure donkey anti-rabbit IgG light chain HRP | Jackson Immuno-Research | #211-032-171 | 1/1000 – 1/40,000 | WB |
| AffiniPure goat anti-mouse  IgG HRP | Jackson Immuno-Research | #715-035-150 | 1/1000 – 1/40,000 | WB |
| 4-HNE | Abcam | ab46545 | 1/625 | IHC |
| LY-6g AlexaFluor488 | Biolegend | Clone 1A8  Cat#127626 | 2.3μg/mouse | IVM |
| CD31/PECAM-1 PE | eBioscience, Invitrogen, Thermo Fisher Scientific | Clone 390  12-0311-83 | 3.0μg/mouse | IVM |
| F4/80 AlexaFluor750 | AbLab, UBC | Clone BM8 | 2.0μg/mouse | IVM |
| F4/80 PE | eBioscience, Invitrogen, Thermo Fisher Scientific | Clone BM8  12-4801-82 | 1.6μg/mouse | IVM |
| TIM4 AlexaFluor647 | Biolegend | Clone RMT4-54  130022 | 2.5μg/mouse | IVM |
| ATPB AlexaFluor647 | Abcam | ab197649 | 1/100 | FC and IVM |
| Donkey anti-Rat IgG (H+L) Highly Cross-Adsorbed Secondary Antibody, Alexa Fluor 594 | Invitrogen, Thermo Fisher | A-21209 | 1/200  (10ug/mL) | IF |
| **Chemicals, reagents, & materials** |  |  |  |  |
| NHS ester AlexaFluor647 | Invitrogen, Thermo Fisher Scientific | A20006 | 25ng/mL | IVM |
| Subtilisin A (Protease from Bacillus licheniformis) type VIII | Sigma-Aldritch | P5380 | 4mg/mL  0.2mg/mL | Mito isolation |
| JC-1 | Invitrogen, Thermo Fisher Scientific | T3168 | 2uM | FC |
| MitoTracker DEEP RED | Invitrogen, Thermo Fisher Scientific | M22426 | 100nM | FC and IVM |
| Syto60 | Invitrogen, Thermo Fisher Scientific | S11342 | 10uM | IVM |
| MitoTracker RED | Invitrogen, Thermo Fisher Scientific | M22425 | 100nM | IVM |
| MitoTracker GREEN | Invitrogen, Thermo Fisher Scientific | M7514 | 200nM | IVM |
| Protein Block | DAKO | X090930-2 | Undiluted | IHC |
| Hydrogen peroxide | Fischer Chemical | H325500 | 3% | IHC |
| IHC secondary antibody | DAKO | K4003 | Undiluted | IHC |
| Antimycin A | Sigma-Aldritch | A8674 | 50uM | Mouse study |
| Diphtheria toxin | Sigma-Aldritch | D0564 | 10ng/g | Mouse study |
| Clodronate and PBS liposomes | LIPOSOMA | CP-005-005 | 50ug/g | Mouse study |
| 700-900nm JADE GREEN microparticles | SpheroTech | FP-0878-2 | 1/30 | FC |
| 1700-2200nm NILE RED microparticles | SpheroTech | FP-2056-2 | 1/30 | FC |
| 1-4μm polyethylene beads | Cospheric | CPMS-0.96 1-4um - 0.2g | 8x10^7^/mouse | Mouse study |
| FluoSpheres™ Sulfate-Modified Microspheres, 00.3.02–4.0μm | Thermo Fisher Scientific | F8845 | 0.002% (w/v) | IVM |
| 5µm nylon filters | PluriSelect | 43-50005-50 | N/A | Mito isolation |
| C-tubes | Miltenyi | 130-093-237 | N/A | Mito isolation |
| Phenol (TE-saturated) | Sigma-Aldritch | 77607 | N/A | ATP in tissue |
| Chloroform | Sigma-Aldritch | C2432 | N/A | ATP in tissue |
| EvaGreen SuperMix | Bio-Rad | 1864034 | N/A | ddPCR |
| EvaGREEN Droplet Oil | Bio-Rad | 1864006 | N/A | ddPCR |
| SYBR GREEN | Applied Biosystems | 4367659 | N/A | qPCR |
| RNeasy Mini Kit | Qiagen | 74104 | N/A | qPCR |
| Qiashredder | Qiagen | 79654 | N/A | qPCR |
| DNA Mini Kit | Qiagen | 51304 | N/A | ddPCR |
| *S. aureus* pH-RODO RED bioparticles | Thermo Fisher | A10010 | 3.5µg/g | IVM |
| pH-RODO RED | Thermo Fisher | P36600 | 25ng/mL | IVM and FC |
| Colibri Retractors | Fine Science Tools | 17000-03 | N/A | Mouse surgery |
| Retractor - 2 Pronged Blunt | Fine Science Tools | 17023-13 | N/A | Mouse surgery |
| ZEISS S100/OPMI PICO | CARL ZEISS CANADA LTD | N/A | N/A | Mouse surgery |
| Vannas Spring Scissors - 3mm Cutting Edge | Fine Science Tools | 15000-10 | N/A | Mouse surgery |
| S&T Vascular Clamps | Fine Science Tools | 00396-01 | N/A | Mouse surgery |
| Schwartz Micro Serrefines | Fine Science Tools | 18052-01 | N/A | Mouse surgery |
| **ELISAs and commercial assays** |  |  |  |  |
| ATP determination kit | Thermo Fisher | A22066 | N/A | Mito isolation |
| Luminescent ATP Detection Assay Kit | Abcam | ab113849 | N/A | ATP in tissue |
| Mouse IL-6 Quantikine ELISA | R&D systems | M6000B | N/A | ELISA |
| Mouse TNFα Quantikine ELISA | R&D systems | MTA00B | N/A | ELISA |
| Mouse CCL2 ELISA | R&D systems | SPCKC-MP-007174 V5 | N/A | ELISA |
| Mouse HGF ELISA | R&D systems | MHG00 | N/A | ELISA |
| Mouse IL-10 Quantikine ELISA | R&D systems | M1000B | N/A | ELISA |
| Mouse 8-isoprostane ELISA | Cayman Chemicals | 516351 | N/A | ELISA |
| **Software** |  |  |  |  |
| Prism (GraphPad) | Unity Health Toronto | v10.1.1 | N/A | Statistical analysis |
| HALO | KRCBS | v2.3.2089.23 | N/A | Histology analysis |
| ZEN | KRCBS | Blue 2012 | N/A | Histology analysis |
| IMARIS | KRCBS | 8.0.2 & 9.6.0 | N/A | IVM analysis |
| **Equipment** |  |  |  |  |
| Automatic tissue processor | Leica | TP1020 | N/A | Histology |
| Paraffin embedder | Leica | EG1160 | N/A | Histology |
| Microtome | Leica | RM2235 | N/A | Histology |
| Autostainer | Leica | Autostainer XL | N/A | Histology |
| Thermal Cautery Unit Ea. | Geiger Instruments Co. 150ST | #CIA1036466 | N/A | IVM |

**Table S2. Mouse-specific primers for ddPCR, qPCR and genotyping.**

| **Target** | **Sequences** | **Purpose(s)** |
| --- | --- | --- |
| **mt-ND1** | Sense: 5’-GCTTTACGAGCCGTAGCCCA-3’  Antisense: 5’-GGGTCAGGCTGGCAGAAGTAA-3’ | ddPCR |
| **mt-ND4** | Sense: 5’-CGCCTACTCCTCAGTTAGCCA-3’  Antisense: 5’-TGATGTGAGGCCATGTGCGA-3’ | ddPCR |
| **B-actin** | Sense: 5’-ATATCGCTGCGCTGGTCGTC-3’  Antisense: 5’-ATAGGAGTCCTTCTGACCCATT-3’ | ddPCR and qPCR |
| **B2M** | Sense: 5'-GACCAAGACTCGTGAGGATAAC-3'  Antisense: 5'-CCAGTGTTGGGTCAGGTTTA-3' | ddPCR and qPCR |
| **GAPDH** | Sense: 5′-AGAAACCTGCCAAGTATGATGACA-3′  Antisense: 5′TGAAGTCGCAGGAGACAACCT-3′ | qPCR |
| **TNFα** | Sense: 5′-TATGGCTCAGGGTCCAACTC-3′  Antisense: 5′-CTCCCTTTGCAGAACTCAGG-3′ | qPCR |
| **IL-10** | Sense: 5′-GGCGCTGTCATCGATTTCTC-3′  Antisense: 5′-GCCTTGTAGACACCTTGGTCTTG-3′ | qPCR |
| **IL-6** | Sense: 5’-GTCCTTCCTACCCCAATTTCCA-3’  Antisense: 5’-TGGTCTTGGTCCTTAGCCAC-3’ | qPCR |
| **CRIg knockout** | Sense common: 5’-CCACTGGTCCCAGAGAAAGT-3’  Antisense wild type-specific: 5'-CACTATTAGGTGGCCCAGGA-3'  Antisense knockout-specific: 5'-GGGAGGATTGGGAAGACAAT-3' | Genotyping |
| **DTR-Lox** | Sense common: 5’-AAAGTCGCTCTGAGTTGTTAT-3’  Antisense wild type-specific: 5'-GGAGCGGGAGAAATGGATATG-3 '  Antisense knockout-specific: 5'-GGCGAAGAGTTTGTCCTCAACC-3' | Genotyping |
| **Clec4f-Cre** | Sense common: 5’- CAAGAAGTCCACAGGGTGGT-3’  Antisense wild type-specific: 5'-GAAAGACCCAAGGGAAGGAG-3 '  Antisense knockout-specific: 5'-ACACCGGCCTTATTCCAAG-3' | Genotyping |
