## Supplementary figures and images for "Mitochondrial transplantation: A novel therapy for liver ischemia/reperfusion injury"

### Supplementary figures 1 and 2

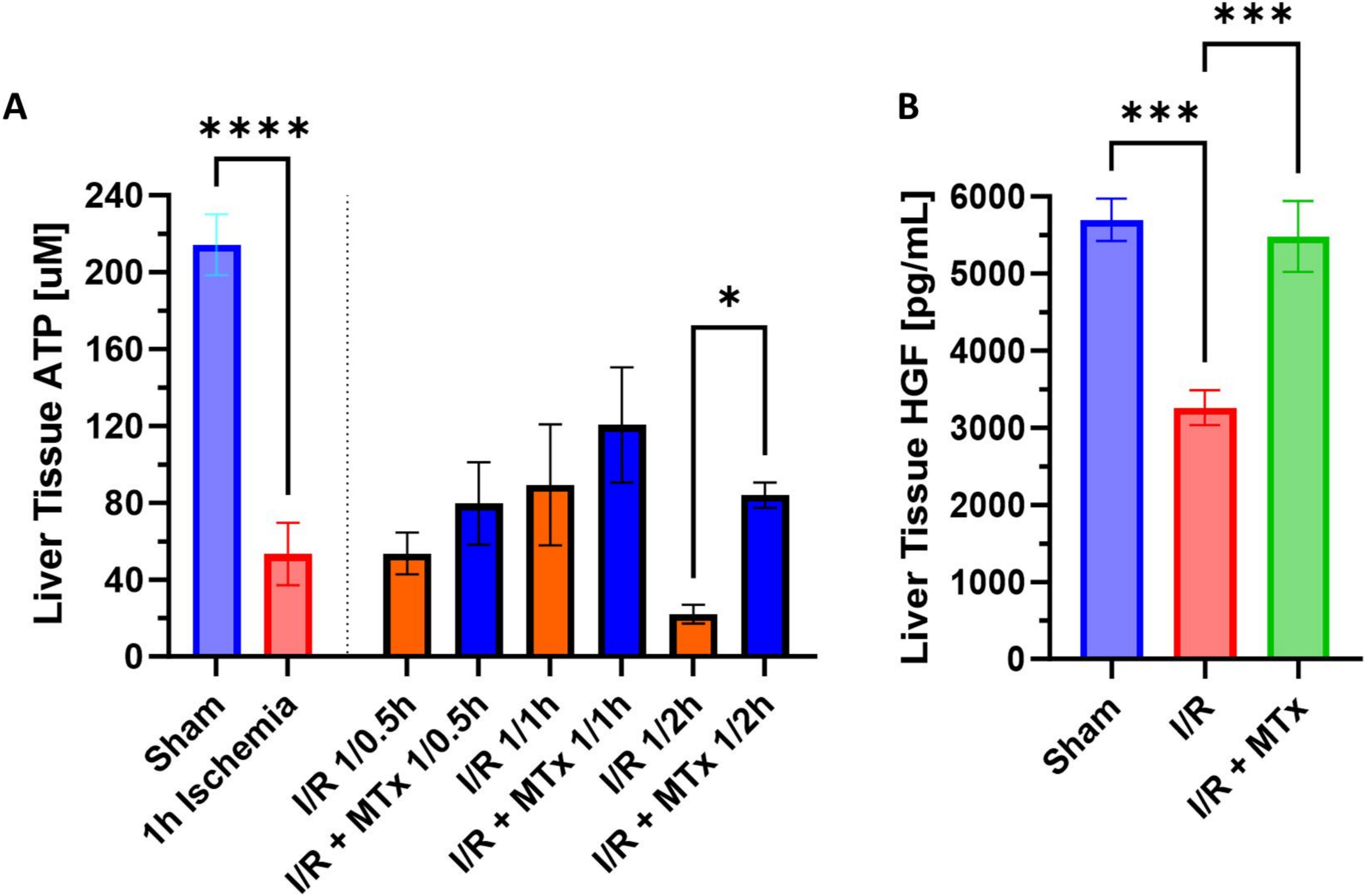

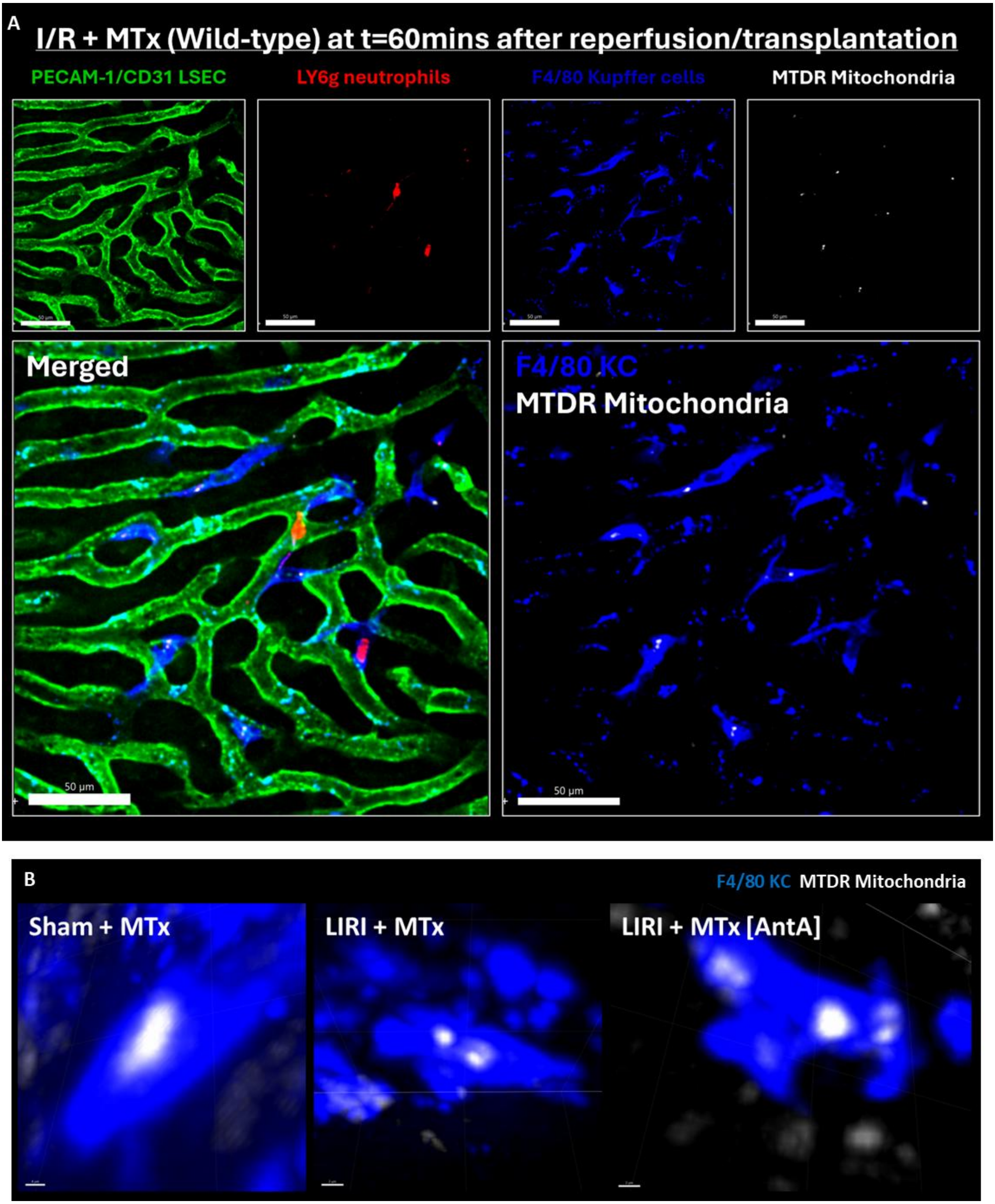
